# SPECK: An Unsupervised Learning Approach for Cell Surface Receptor Abundance Estimation for Single Cell RNA-Sequencing Data

**DOI:** 10.1101/2022.10.08.511197

**Authors:** Azka Javaid, H. Robert Frost

## Abstract

The rapid development of single cell transcriptomics has revolutionized the study of complex tissues. Single cell RNA-sequencing (scRNA-seq) can profile tens-of-thousands of dissociated cells from a tissue sample, enabling researchers to identify cell types, phenotypes and interactions that control tissue structure and function. A key requirement of these applications is the accurate estimation of cell surface protein abundance. Although technologies to directly quantify surface proteins are available, this data is uncommon and limited to proteins with available antibodies. While supervised methods that are trained on Cellular Indexing of Transcriptomes and Epitopes by Sequencing (CITE-seq) data can provide the best performance, this training data is also limited by available antibodies and may not exist for the tissue under investigation. In the absence of protein measurements, researchers must estimate receptor abundance from scRNA-seq data. We thereby developed a new unsupervised method for receptor abundance estimation using scRNA-seq data called SPECK (Surface Protein abundance Estimation using CKmeans-based clustered thresholding) and evaluated its performance against other unsupervised approaches on up to 215 human receptors and multiple tissue types. This analysis reveals that techniques based on a thresholded reduced rank reconstruction (RRR) of scRNA-seq data are effective for receptor abundance estimation with SPECK providing the best overall performance.

## Introduction

Transcriptome profiling has conventionally been performed using RNA sequencing (RNA-seq) of bulk tissue samples (1, 2). While bulk RNA-seq facilitates genome-wide gene expression profiling, it measures the average gene expression for all cells within a tissue sample. To address the limitations of conventional RNA-seq, single cell RNA-sequencing (scRNA-seq) technologies (3), such as the 10X Chromium System (4), have been developed that can quantify gene expression within thousands of cells from a single tissue sample. The resolution provided by single cell assays is critical for unraveling the biology of diseases associated with complex tissues, e.g., the tumor microenvironment, with important implications for advancing precision medicine, e.g., understanding the association between specific driver mutations and the composition and phenotype of tumor infiltrating immune cells. Key applications of scRNA-seq data include cataloging the cell types in a tissue (5), reconstructing dynamic processes (6), and analyzing cell-cell signaling (7). Many of these tasks are based on the abundance of cell surface proteins, which follows the traditional use of immunohistochemical profiling for characterizing dissociated cells (e.g., Fluorescence Activated Cell Sorting (FACS) (8)) or intact tissue. Although techniques such as Cellular Indexing of Transcriptomes and Epitopes by Sequencing (CITE-seq) (9) can measure protein abundance for individual cells using barcoded antibodies, this type of data is uncommon, and targets a limited number of proteins that have available antibodies.

If receptor protein measurements are not available for a given sample, researchers can estimate receptor abundance from scRNA-seq data using supervised or unsupervised approaches. Supervised methods, such as the single cell Transcriptome to Protein prediction with deep neural network (cTPnet) tool (10), fit a predictive model on training data that captures both gene expression and protein abundance, e.g., joint scRNA-seq/CITE-seq, and use the trained model to generate estimates for target scRNA-seq datasets. While supervised methods can provide excellent predictive performance, they are only feasible if sufficient training data exists and may be susceptible to over-fitting and low transparency. For example, cTPnet only supports 24 immune-related receptors and uses a deep learning approach that makes model interpretation challenging. For cases where training data is not available, which includes many receptors that lack CITE-seq antibodies, an unsupervised approach that leverages RNA expression to determine protein levels must be employed. While the heterogeneity induced by processes such as transcriptional bursting (11) may reduce the otherwise 56%-84% reported protein variation accounted for by RNA expression measured using canonical quantification tools (12), advances in single cell technology may make it increasingly more feasible to use RNA expression to quantify protein abundance. Abundance estimation using gene expression data is further legitimized in the context of relative abundance estimation. While post-transcriptional regulation has a substantial influence on absolute protein concentration, these processes do not have a major impact on relative protein levels (13).

One simple unsupervised strategy uses the expression of the associated RNA transcript as a proxy for protein abundance. While this approach can work in some situations, the sparsity of scRNA-seq data often leads to poor quality estimates. Although this sparsity can be mitigated by a cluster-based approach that approximates abundance using the average expression across all cells in a cluster, such a cluster-level analysis is sensitive to the number of computed clusters, and, importantly, ignores within cluster heterogeneity. A second promising family of unsupervised methods for generating cell-specific estimates are scRNA-seq data imputation techniques. In this scenario, the imputed value of the receptor transcript is used to estimate protein abundance under the assumption that the imputation process reduces sparsity without inflating false positives. Although a large number of scRNA-seq imputation approaches exist (14), we have found that the class of reduced rank reconstruction (RRR) techniques, which assumes that the intrinsic dimensionality of scRNA-seq data is much lower than the empirical rank, provide superior performance relative to other methods. RRR-based imputation methods include Markov Affinity-based Graph Imputation of Cells (MAGIC) (15) and Adaptively thresholded Low-Rank Approximation (ALRA) (16). While MAGIC and ALRA significantly outperform the naïve approach of directly using the receptor transcript, they have limitations. ALRA, for example, applies the same quantile probability of 0.001 to threshold each gene, thereby overlooking considerable individual, gene-level expression variability. While MAGIC, like ALRA, is scalable, its performance is largely inferior to ALRA’s as a single cell imputation strategy (16).

To address the limitations of existing unsupervised approaches for estimating relative receptor protein abundance, we developed a new technique named Surface Protein abundance Estimation using CKmeans-based clustered thresholding (SPECK). Similar to ALRA, the SPECK method utilizes a singular value decomposition (SVD)-based RRR but includes a novel approach for thresholding of the reconstructed gene expression matrix that improves receptor abundance estimation. A second important contribution of this paper is a comprehensive evaluation of unsupervised receptor abundance estimation performance across 215 human cell surface receptors and multiple tissue types.

## Methods

### SPECK method for scRNA-seq data

The SPECK method estimates abundance profiles for *m* cells and *n* genes in a *m × n* matrix of scRNA-seq counts using the procedure detailed below. See Figure 1 and Algorithm 1 for the implementation overview and the associated pseudocode, respectively.

**Fig. 1.**
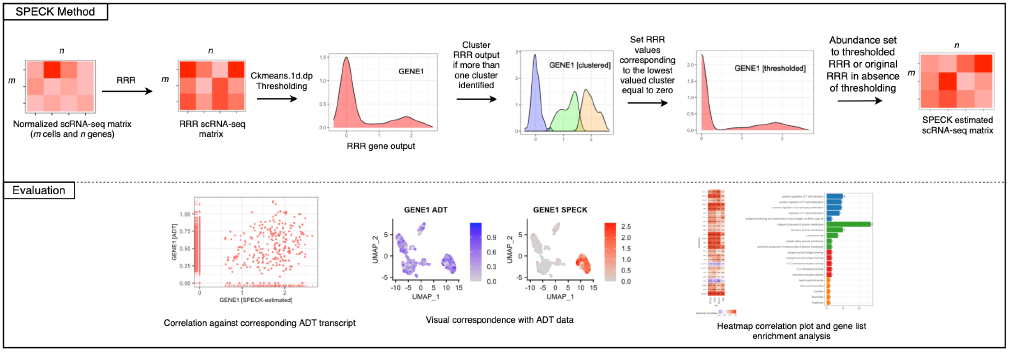
SPECK Method: SPECK performs normalization, rank selection, reduced rank reconstruction and thresholding on a *m × n* scRNA-seq count matrix with *m* cells and *n* genes. The reconstructed and thresholded matrix is of size *m × n*. To evaluate SPECK, receptor abundance estimates were visually assessed using feature plots and heat maps and correspondence to CITE-seq data was quantified using the Spearman rank correlation.

### Rank estimation and reduced rank reconstruction (RRR)

Given a *X_m,n_* scRNA-seq count matrix, whose elements represent the number of mRNA molecules associated with each gene detected in a specific cell, we first performed log-normalization using Seurat’s normalization pipeline (17) to generate relative expression values. Seurat’s log-normalization procedure divides the gene-level counts for each cell by the total counts for all genes in that cell, multiplies this value by a scale factor of 10,000 and then natural-log transforms the product. A singular value decomposition (SVD)-based reduced rank reconstruction (RRR) is next performed on the resulting *X_m,n_* normalized matrix to create a low-rank representation of the original gene expression data. Because genes act in a concerted and interdependent manner, the original expression values are well approximated by a low-rank matrix produced by the RRR procedure (18, 19). To perform the RRR, we leveraged the randomized SVD algorithm implemented in the rsvd R package (20), which was observed to be accurate and computationally efficient relative to non-randomized truncated SVD techniques. This randomized SVD method was used to generate a rank-100 decomposition of the *X_m,n_* normalized expression matrix. To estimate the exact rank for the final RRR, the singular values from this rank-100 SVD decomposition were used to compute the standard deviation of the non-centered sample principal components and a rate of change in these standard deviations between successive components was calculated. The target rank *k* was then selected to represent a numeric value, ranging from 1 to 100, for which the absolute value of the difference between consecutive standard deviation estimates was at least 0.01 for two or more estimate pairs. Although this rank selection procedure is a heuristic method, motivated by the commonly used elbow method for determining the number of principal components, we find that it works well in practice. The first *k* columns from each of the three rank-100 rsvd-based decomposition matrices were subsequently used to generate a rank *k* reconstruction of the *X_m,n_* normalized expression matrix.

### Cluster-based thresholding

We next performed clustered thresholding on the RRR values for each of the *n* genes from the *X_m,n_* RRR matrix. This step was inspired by the bimodality often exhibited by protein expression distributions, which functionally reflects the presence of two different sub-populations with varying individual response mechanisms to factors like stress or a particular drug treatment (21, 22). While bimodality in protein expression at the population level has historically been linked to stochastic switching (23), defined by the transition of cells between multiple phenotypes as a result of environmental fluctuations, it is now more recently attributed to cell-to-cell variability that affects the sustained oscillation frequencies in a heterogeneous cell population (21). We sought to leverage this idea of population-level bimodality exhibited by proteomics data in our abundance estimation technique by performing a thresholding step on the RRR values to ensure that the resulting estimates were representative of the corresponding CITE-seq data.

For this purpose, we utilized a one-dimensional (1-D) clustering algorithm from the Ckmeans.1d.dp package (24, 25) to perform thresholding on the reconstructed expression values for each gene. This algorithm functions by reducing the overarching problem of clustering a 1-D array consisting of *x*_1_*,…, x_n_* values into *k* clusters to a sub-problem of minimizing the sum of squares of within-cluster distances from an element to its associated cluster mean for clustering *x*_1_*,…, x_i_* values into *m* clusters. Dynamic programming is then used to solve this recurrence equation and find the assignment of all *n* values to *k* clusters. As noted, this iterative computation reduces the time complexity of the algorithm to *O*(*nk*), which compares with *O*(*qknp*) for standard k-means algorithm, where *q* defines the number of iterations and *p* specifies the dimensionality (25, 26). We selected this dynamic programming-based 1-D clustering algorithm since in addition to its runtime efficiency, it is optimal and, unlike comparative methods like k-means, is not dependent on the definition of initial cluster centers, thereby producing consistent cluster assignments for each run. The Ckmeans.1d.dp algorithm implementation requires specification of *k*, which defines the minimum and maximum number of clusters to be examined. While, given the bimodal expression pattern of proteomics data, an upper bound of two seems reasonable for this Ckmeans.1d.dp-based parameter *k*, we specified a slightly higher upper bound to account for potential additional expression multimodality in CITE-seq data beyond that accounted for by bimodality. We found that the upper limit of four was appropriate to ensure that cells were mapped to a fairly granular and interpretable number of clusters. Further analysis revealed that the thresholding performance was relatively insensitive to this upper bound, as specified within the range of 3-15. With the minimum cluster number set to one and the maximum cluster number set to four, we performed thresholding only if more than one cluster was identified. If so, then all the non-zero values in the RRR output for the gene corresponding to the indices of the least-valued cluster, as identified by the cluster mean, were set to zero. The zero values corresponding to the indices of the least-valued cluster and all other non-zero and zero values corresponding to higher-valued clusters in the RRR gene output were preserved. If only one cluster was identified, thresholding was not performed and the RRR gene values were preserved. The final output consisted of a *X_m,n_* thresholded RRR matrix.

## Evaluation

### Comparison methods

Comparative evaluation of the SPECK method was performed against three other approaches described below.

- ALRA: Receptor abundance was estimated using the imputed value of the associated transcript generated by the ALRA method with v.1.0.0 (16). scRNA-seq data was normalized using ALRA’s procedure, which normalizes each cell column by a factor of 10,000 followed by adding a pseudocount of one and taking the logarithm of each entry. Log-normalized data was then RRR and imputed using ALRA’s rank-estimation approach, which chooses the largest value of *k*, given an upper bound of *k* < 100, such that the gap between consecutive singular values *s_k_* is significantly different from the mean and standard deviation of typical noise spacings, defined to be *s*_80_ *… s*_100_ (i.e., *s_k_ > μ* + 6*σ*).
- MAGIC: Receptor abundance was estimated using the imputed value of the associated transcript generated by the MAGIC method with Rmagic v.2.0.3 (15). scRNA-seq data was normalized using the library size (i.e., transcript abundances) to ensure that each cell has the same transcript count. Normalized data was subsequently imputed with the number of nearest neighbors parameter *knn* set to 15.
- RNA transcript: Receptor abundance was estimated using the normalized value of the associated transcript with the normalization performed using Seurat’s log-normalization procedure (17).

**Algorithm 1.**
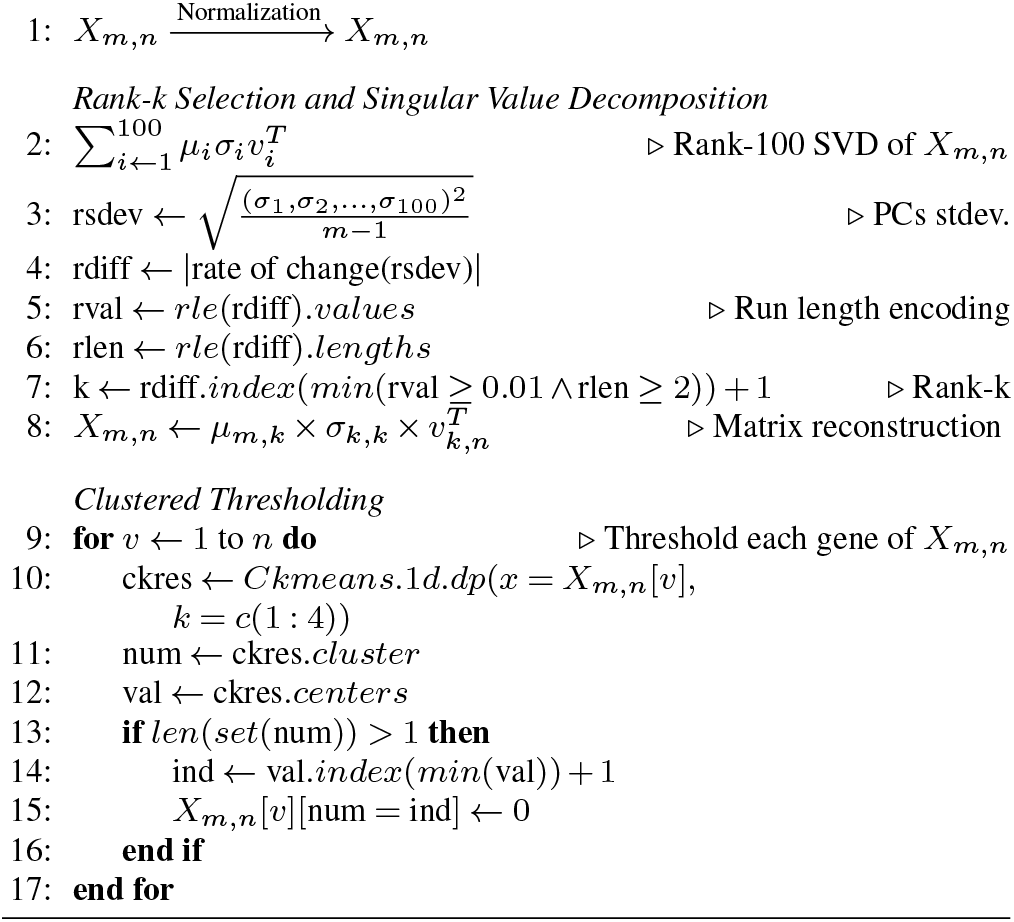
SPECK Algorithm.

### Public single cell data

SPECK was evaluated relative to the ALRA, MAGIC and RNA transcript methods on two publicly accessible joint CITE-seq/scRNA-seq datasets: 1) the Hao et al. (27) human peripheral blood mononuclear cell (PBMC) dataset (GEO (28) series GSE164378) that contains 161,764 cells profiled using 10X Chromium 3’ with 228 TotalSeq A antibodies, and 2) the Stuart et al. (29) human bone marrow mononuclear cell (BMMC) dataset (GSE128639) that contains 33,454 cells similarly profiled with 25 TotalSeq A antibodies. For both datasets, the scRNA-seq data was used to generate receptor abundance estimates that were evaluated relative to the corresponding CITE-seq ADT data. Each dataset was individually processed using Seurat v.4.1.0 (17, 27, 29, 30) in R v.4.1.2 (31). CITE-seq ADT counts were normalized using the Seurat implementation of the centered log-ratio transformation (32).

### Setup and evaluation metrics

We evaluated the SPECK-estimated abundance profiles on two different joint scRNA-seq/CITE-seq datasets detailed above: a PBMC dataset with expression measurements for 33,538 genes and CITE-seq measurements for 228 receptor antibodies and a BMMC dataset with expression measurements for 17,009 genes and CITE-seq measurements for 25 receptor antibodies. To explore both the variability in estimation performance and how the number of cells impacts performance, five random subsets were generated from each dataset for different numbers of cells with subset size varying between 1000 and 60,000 cells for the PBMC data and between 1000 and 30,000 cells for the BMMC data. We determined the upper limit of 60,000 cells for the PBMC data by the ability to perform RRR on 16 CPU cores without any virtual memory allocation errors. From the initial 228 antibodies included in the PBMC data, antibodies mapping to multiple HUGO Gene Nomenclature Committee (HGNC) (33) symbols were removed. Since assessment was performed for a varying number of cells, receptors not expressed in smaller cell groups were dropped. Our final assessed receptors for the PBMC data ranged from 205 to 215. For each random subset, we measured the correspondence between the estimated receptor abundances and the CITE-seq ADT measurements using the Spearman rank correlation. Spearman correlation was chosen since it can quantify relationships between the ranks of two variables, as opposed to their absolute values, which is appropriate for comparing the estimated relative abundance profiles produced by the SPECK, ALRA, MAGIC and the RNA transcript methods against CITE-seq data. The Spearman correlation metric is also robust to outliers and can be used to analyze nonlinear relationships (34). SPECK’s performance, as quantified by the average Spearman rank correlation coefficient across the five random subsets for a given number of cells, was compared with the performance of imputed expression profiles generated by ALRA and MAGIC and the normalized RNA transcript counts.

## Results

### SPECK improves correspondence of abundance estimates with CITE-seq ADT data

We first quantified the proportion of receptors where the estimated abundance generated by either the SPECK, ALRA, MAGIC or RNA transcript method had the highest average rank correlation values with CITE-seq ADT values (Figures 2 and 3). To examine the specific impact of SPECK’s thresholding procedure, we also quantified proportion of receptors for which either the complete SPECK method (i.e., both RRR and thresholding) or the SPECK method without thresholding (i.e., just RRR) generates the best relative estimate (Figures 4 and 5). The statistical significance of the individual difference between the proportion of receptors for which SPECK had the best average correlation and the proportion for the method with the next best average correlation and the difference between the proportion of receptors for which the complete SPECK method had the best average correlation and the proportion for the SPECK method without thresholding with the best average correlation were quantified using a two-sample z-test of proportions. The collection of p-values from all comparisons were treated as a family of hypotheses. The Benjamini–Hochberg method (35) was used to control the False Discovery Rate (FDR) for multiple comparisons.

**Fig. 2.**
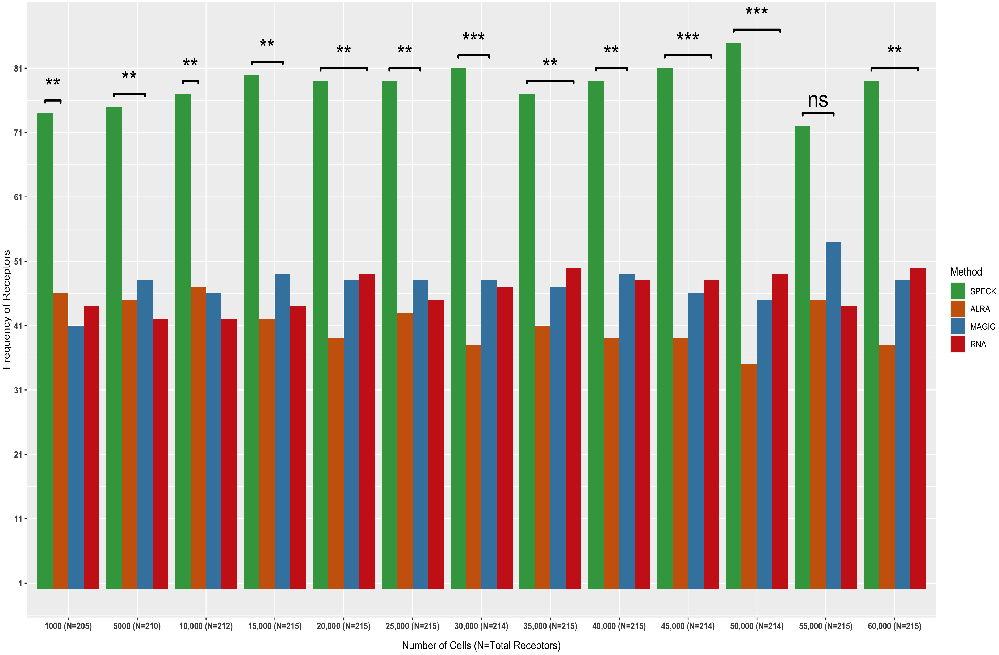
Frequency of receptors with the highest averaged rank correlation values between PBMC ADT data and abundance estimates produced by SPECK or imputed profiles produced by ALRA, MAGIC or RNA for five subsets of cells ranging from 1000 to 60,000 for the PBMC data are displayed. Asterisks indicate the adjusted p-value significance levels for differences between the top two, pairwise proportions for each cell category (*\*\*\**, 0.0001 < *P* < 0.001; *\*\**, 0.001 < *P* < 0.01; *\**, 0.01 < *P* < 0.05;ns, 0.05 < *P* < 1).

**Fig. 3.**
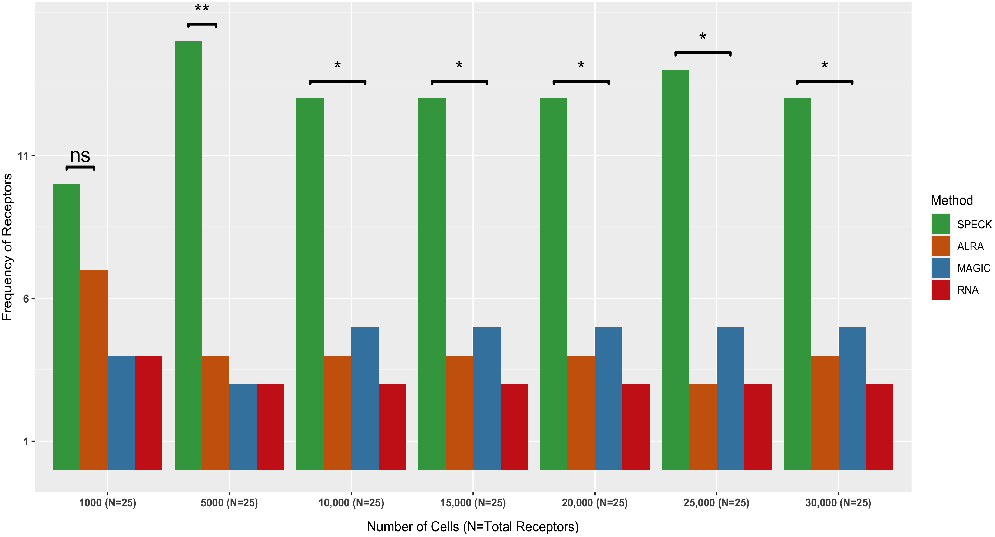
Frequency of receptors with the highest rank correlation between BMMC ADT data and abundance profiles produced by a specified estimation/imputation technique, including SPECK, ALRA, MAGIC and RNA, averaged over five subsets of cells ranging from 1000 to 30,000, are indicated. Asterisks indicate the adjusted p-value significance levels for differences between the top two proportion values (*\*\**, 0.001 < *P* < 0.01; *\**, 0.01 < *P* < 0.05;ns, 0.05 < *P* < 1).

**Fig. 4.**
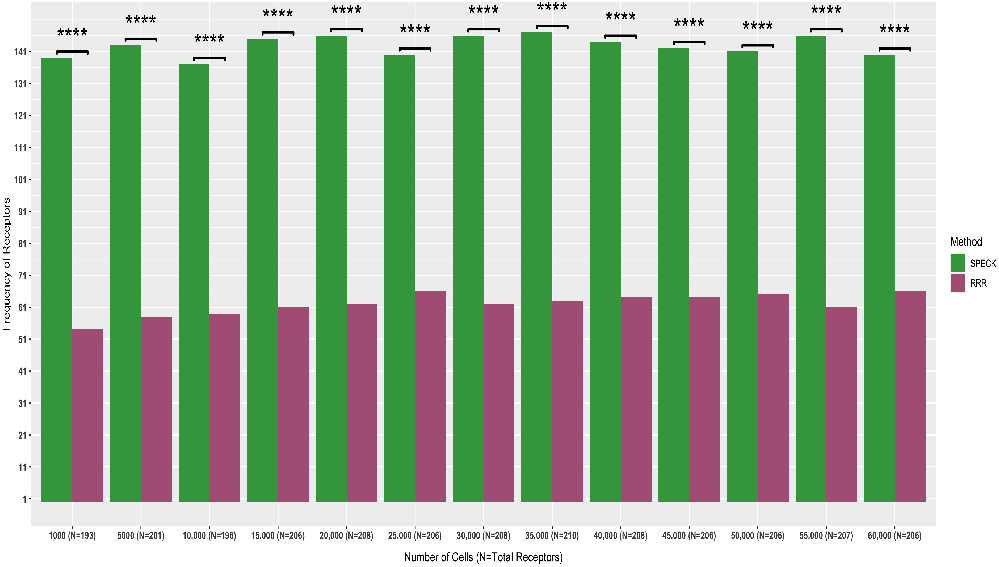
SPECK-based RRR and thresholded estimates are compared with SPECK-based RRR-only values and frequency of receptors with the highest Spearman rank correlation between PBMC ADT data and estimated values produced by either estimates, averaged over five subsets of cells ranging from 1000 to 60,000, are displayed. Receptors estimated with both techniques that have the same correlation value against corresponding ADT data are removed. Asterisks indicate the significance levels of p-values from the proportion tests (*\* * \*\**, 0 < *P* < 0.0001).

**Fig. 5.**
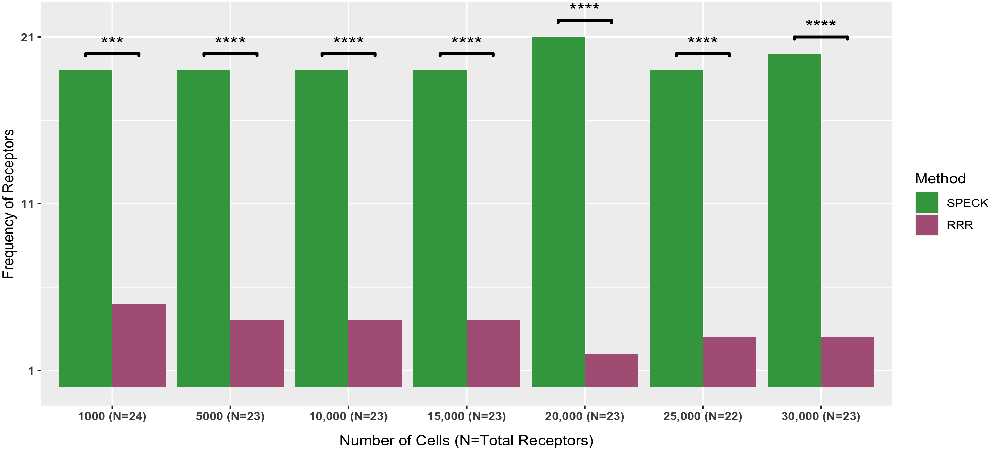
Frequency of receptors with the highest rank correlation between BMMC ADT data and abundance profiles produced by SPECK (RRR and thresholding) and SPECK-based RRR-only, as averaged over five subsets of cells ranging from 1000 to 30,000, are shown. Receptors with ties in rank correlation values are removed. Asterisks indicate the p-value significance levels for the proportion tests (*\* * \*\**, 0 < *P* < 0.0001; *\* * \**, 0.0001 < *P* < 0.001).

As Figures 2 and 3 show, SPECK estimated abundance profiles consistently have a better correspondence with CITE-seq ADT data compared to the comparative methods across all data subsets for the PBMC and BMMC datasets. For example, for the 10,000 cell subsets from the PBMC data, the SPECK estimated abundance values have the highest correlation with CITE-seq ADT values for 76 receptors (36%). This compares with 47 (22%) receptors where ALRA generates the best estimates, 46 (22%) receptors where MAGIC generates the best estimates, and 42 (20%) receptors where the normalized RNA transcript abundance provides the best estimate. The difference between the proportion of receptors where SPECK provides the best estimate and the proportion for the next best method is statistically significant at a FDR level of 0.05 for all subset sizes except the 55,000 cell category for the PBMC data and the 1000 cell category for the BMMC data. We also observed that SPECK’s performance is relatively insensitive to cell subset size and is numerically better than comparative imputation abundance estimates across all subsets. Figure 4 indicates the full SPECK method generates superior estimates relative to the no thresholding version of SPECK on the PBMC data. Comparable results were found for the BMMC data, as shown by Figure 5. All differences in receptor proportions were statistically significant at a FDR level of 0.05 for every subset size for the PBMC and the BMMC datasets. These results emphasize the central contribution of SPECK’s thresholding procedure in considerably improving the accuracy of the estimated abundance profiles over an RRR-only approach.

### Examining individual abundance values using density estimation

We next visually assessed correspondence between the estimated abundance profiles produced by SPECK, ALRA, MAGIC and the RNA transcript method with analogous CITE-seq ADT data using kernel density estimation-based smoothed scatter plots for random subsets of 10,000 cells from the PBMC and the BMMC datasets. This subset size was selected since 10,000 cells present a reasonable cut off point between a small-sized and a comparatively large scRNA-seq dataset. The CD14, CD19 and CD79b receptors were chosen for visualization since each receptor functions as an important marker for an immune cell population. While CD14 is a marker for monocytes, types of leukocytes that actively contribute to an adaptive immune response by differentiating as macrophages and dendritic cells, CD19 and CD79b are markers for B cells, types of lymphocytes that are primarily involved in producing antibodies against foreign antigens and serving as antigen-presenting cells. The lymphocyte to monocyte ratio has further clinical utility as it can serve as an independent predictor of overall survival in patients undergoing curative resection with colorectal cancer (36).

Figures 6 and 7 show these scatter plots for select receptors and, overall, indicate that compared to the abundance profiles generated by ALRA, MAGIC and the RNA transcript method, SPECK estimated abundance profiles for CD14, CD19 and CD79b show a distinctive separation of cells in high-density regions that are highly correlated with CITE-seq data. These results underscore an important contribution of SPECK in producing relatively accurate abundance estimates compared to other unsupervised techniques.

**Fig. 6.**
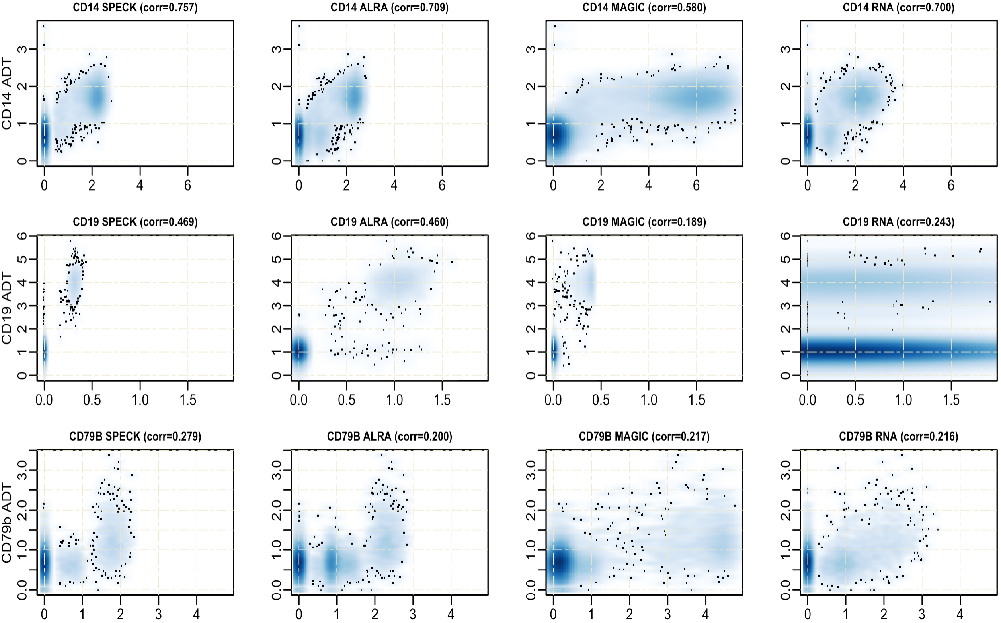
Smoothed scatterplot representation of SPECK-estimated abundance profiles and imputed abundance values produced by ALRA, MAGIC and RNA against ADT data for CD14, CD19 and CD79b receptors. Plot is based on a two-dimensional kernel density estimate of estimated abundance profiles for a subset of 10,000 cells from the PBMC data. The same lower and upper limits for the x-axis scale are set for every receptor based on the range of the estimated abundance profiles produced by SPECK and all other comparative methods.

**Fig. 7.**
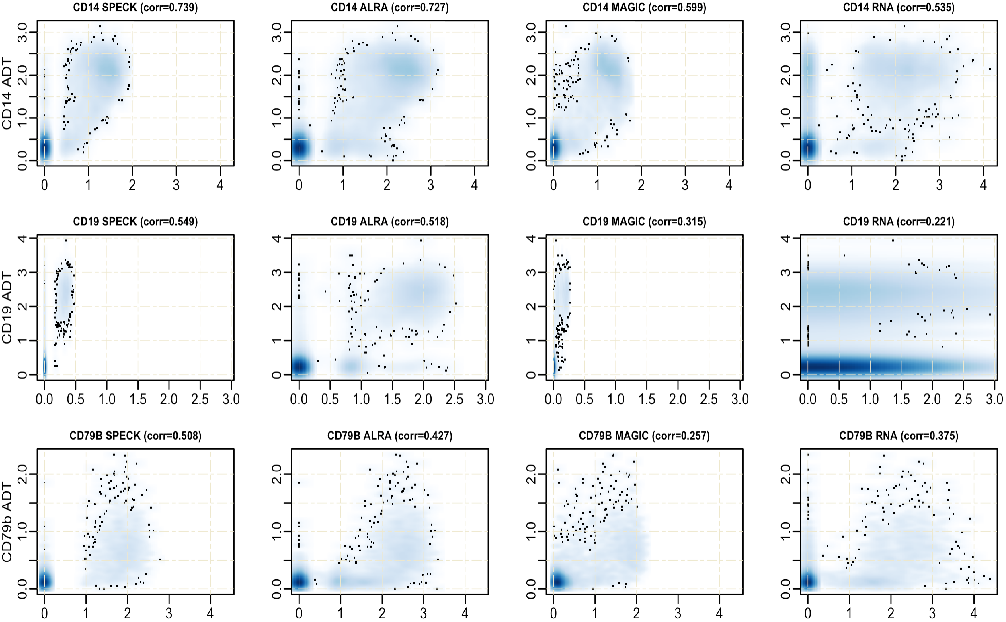
Density-based smoothed scatterplot representation of SPECK-estimated values and ALRA, MAGIC and RNA-imputed abundance profiles against ADT data for CD14, CD19 and CD79b receptors for a subset of 10,000 cells from the BMMC data. The same lower and upper limits of the x-axis are analogously determined based on the range of the abundance estimates produced by all methods combined.

### Visualization of individual estimates using a low-dimensional space

Next, we visually assessed correspondence between the estimated abundance profiles and the CITE-seq ADT data, for the same random subsets of 10,000 cells used for Figures 6 and 7, via a projection of cells onto the first two Uniform Manifold Approximation and Projection (UMAP) (17) dimensions. This second visualization allowed us to capture the estimated abundance profiles on an underlying low-dimensional manifold, thereby enabling mapping of these values over both the local and global topological structure of the transcriptomic data. Figures 8 and 9 show a feature plot visualization of ADT data and the estimated abundance profiles produced by SPECK, ALRA, MAGIC and the RNA transcript techniques for the CD14, CD19 and CD79b receptors from the PBMC and the BMMC datasets, respectively. The abundance estimates produced by SPECK have more visually defined expression profiles that are primarily relegated to region of cells with high CITEseq ADT abundance. This trend is especially visible for the CD79b receptor for both the PBMC and the BMMC datasets, for which ALRA and the RNA transcript techniques especially indicate a homogeneous and less distinctive expression pattern. These results suggest the potential utility of the SPECK method for cell typing of scRNA-seq data.

**Fig. 8.**
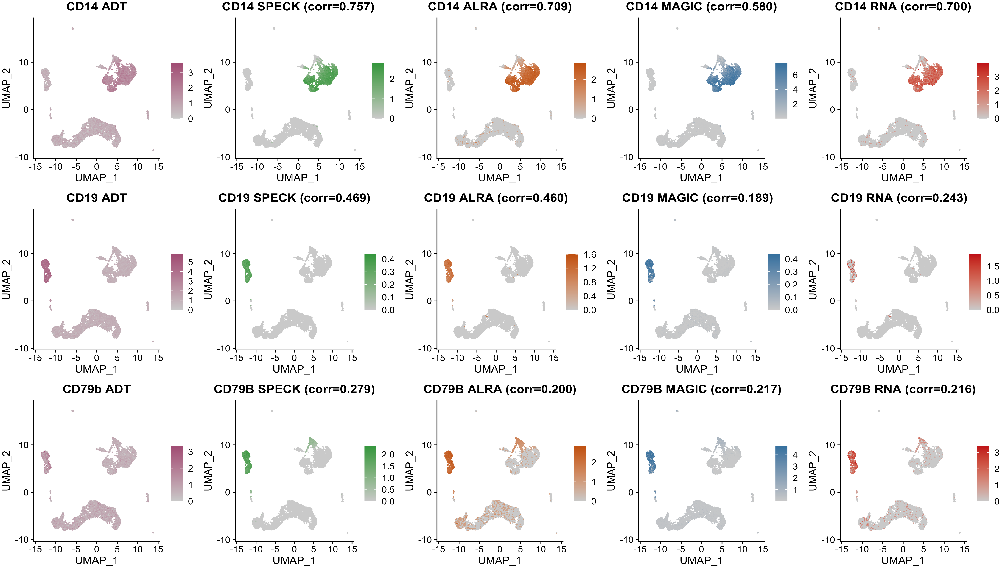
Low-dimensional projection of abundance profiles for CD14, CD19 and CD79b receptors as estimated by SPECK and imputed by ALRA, MAGIC and RNA and corresponding ADT data for a subset of 10,000 cells from the PBMC data.

**Fig. 9.**
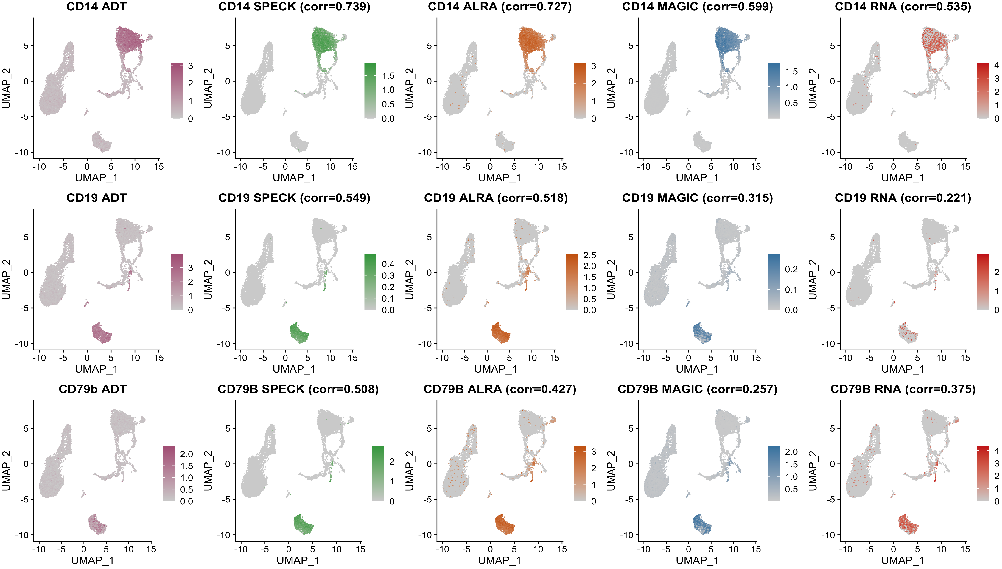
Projection of estimated abundance profiles, produced by SPECK, ALRA, MAGIC and RNA, for CD14, CD19 and CD79b receptors for a subset of 10,000 cells from the BMMC data.

### Framework for selection of an informed abundance estimation strategy

Following the overall evaluation of abundance profiles estimated by SPECK across all subsets and an examination of the distribution of estimated abundance values for select receptors for an individual cell subset, we quantified the magnitude and direction of the rank correlation values between CITE-seq ADT data and the estimated profiles generated by SPECK, ALRA, MAGIC and the RNA transcript method. Figure 10 displays a heatmap plot of these correlation values for 215 receptors averaged over five subsets of 60,000 cells from the PBMC data while Figure 11 shows the heatmap for 25 receptors averaged over five subsets of 30,000 cells from the BMMC data. In addition to confirming the results from Figures 2 and 3, which indicate that SPECK overall outperforms ALRA, MAGIC and the RNA transcript method in the task of abundance estimation, these figures have scientific utility as they can be referenced to determine the most appropriate estimation/imputation strategy for a given receptor. For example, for the CD8a receptor, estimated abundance profiles produced by ALRA are more highly correlated with corresponding ADT data compared to estimated abundance profiles produced by SPECK, MAGIC and RNA for both the PBMC and the BMMC datasets. ALRA may, therefore, be a preferable strategy for estimating abundance profiles for the CD8a receptor compared to other techniques.

**Fig. 10.**
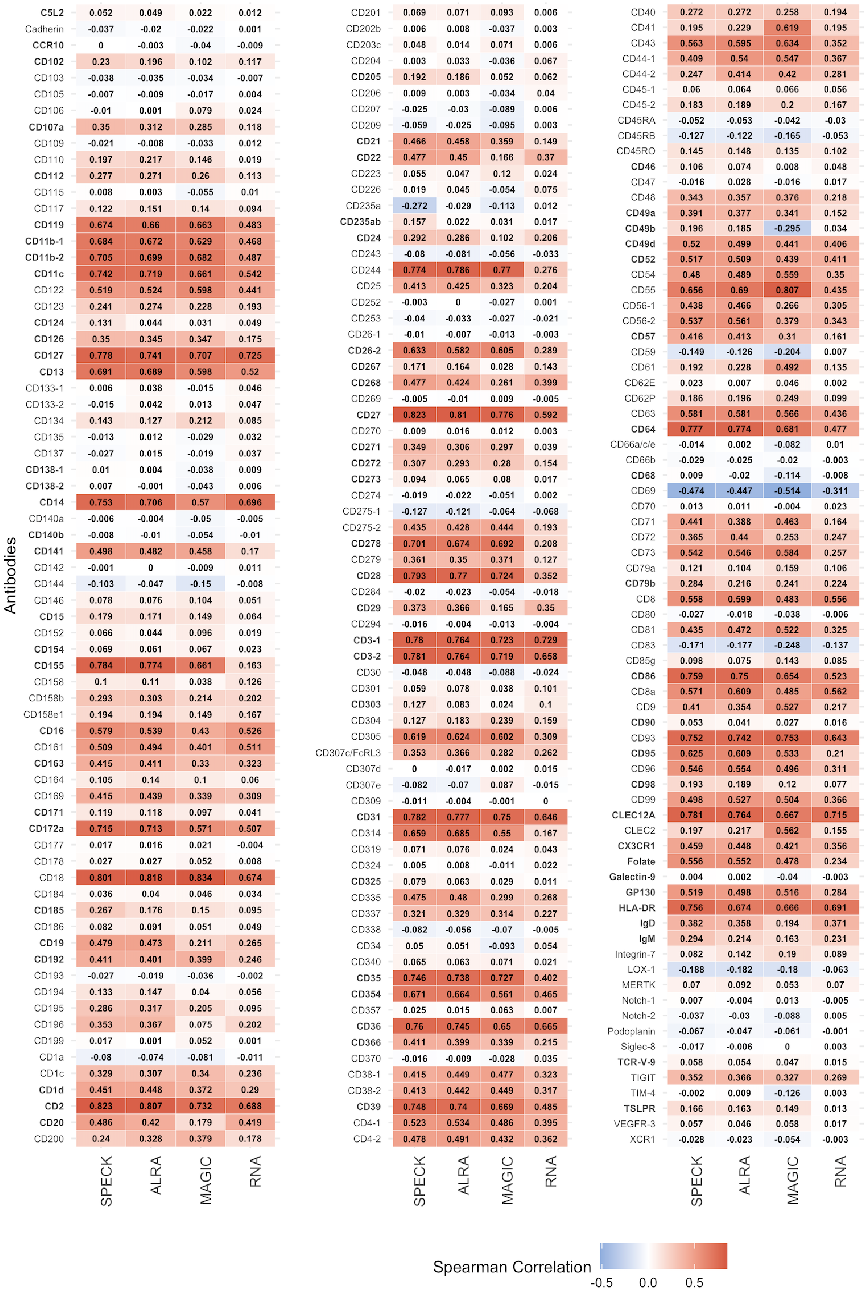
Individual rank correlation values between the PBMC ADT data for 215 receptors and the SPECK-estimated or the ALRA, MAGIC and RNA-imputed abudance values, as averaged over five subsets of 60,000 cells, for 215 receptors are displayed. Bold text format is used to indicate receptors that are estimated with SPECK and are highly correlated with ADT data as compared to imputed profiles produced by ALRA, MAGIC and RNA.

**Fig. 11.**
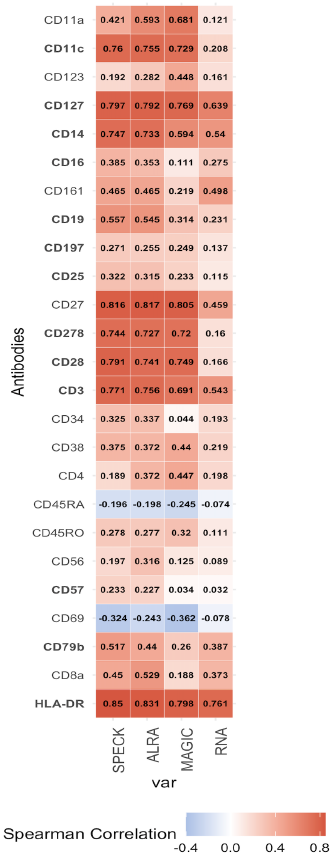
Rank correlation values between the BMMC ADT data and SPECK-estimated abundance profiles or ALRA, MAGIC and RNA-imputed values, as averaged over five subsets of 30,000 cells, for 25 receptors are displayed. Receptors in bold text are estimated with SPECK and are most correlated with ADT data as compared to imputed profiles produced by ALRA, MAGIC and RNA.

## Discussion

In this work, we describe a new unsupervised learning method, SPECK, that uses RRR and thresholding to estimate cell surface receptor abundance from scRNA-seq data. We evaluated our approach on PBMC and BMMC joint scRNA-seq/CITE-seq datasets for a large set of human receptors. This comparative evaluation demonstrates that SPECK generates more accurate receptor abundance estimates over the RRR-based imputation methods ALRA and MAGIC or the direct use of normalized receptor RNA transcript abundance. An important contribution of SPECK is a novel strategy for cluster-based thresholding of the reconstructed gene expression values. Our proposed thresholding mechanism differs from existing thresholding schemes, e.g., the ALRA thresholding approach, in two ways. First, our strategy does not always threshold a gene and, second, does not necessarily apply the same quantile to threshold each gene, thereby enabling a recovery of more accurate estimated abundance profiles. A second important novel contribution of this proposed framework is that it outlines an extensive evaluation strategy for benchmarking the effectiveness of unsupervised methods for estimating receptor abundance using scRNA-seq data. With performance comparisons performed across multiple cell subsets for at least 205 and at most 215 receptors from the PBMC data and 25 receptors from the BMMC data, this evaluation provides information on the relative performance of unsupervised methods across a wide range of human receptors.

One limitation of our proposed approach is that SPECK is not necessarily the best abundance estimation strategy for all receptors. For example, SPECK produces a negative correlation value for CD69 for both the PBMC and the BMMC data as shown by Figures 10 and 11, respectively. While this correlation is in agreement with the negative correlations returned by comparative methods, it still points to a deficiency in the use of transcriptomics data to infer protein abundance. We hope to address this limitation in future work by using pathway analysis to identify the biological characteristics of receptors whose abundance can be accurately estimated from scRNA-seq using unsupervised methods such as SPECK. Such insights could then be leveraged by researchers to help determine whether unsupervised estimation of receptor abundance is feasible for a given investigation or whether direct proteomic measurements are motivated. A second limitation of our approach stems from a key deficiency of scRNA-seq data, namely, that current scRNA-seq protocols can only capture static transcriptional states of cells at a specific point in time. This limitation prevents the applicability of SPECK to surface protein abundance estimation for dynamic biological processes such as tissue regeneration. We aim to address this limitation in future work by providing an implementation of SPECK that can incorporate the recently developed RNA Velocity method (37) for estimating RNA abundance at future transcriptional cell states, thereby extending applicability to abundance estimation for dynamic processes such as cell differentiation. Current R package implementation for SPECK is available on the Comprehensive R Archive Network (CRAN) (38).

In conclusion, SPECK is a promising approach for unsupervised estimation of surface receptor abundance for scRNA-seq data that addresses limitations of existing imputation methods such as ALRA and MAGIC. The cell surface receptor abundance profiles generated by SPECK have important scientific utility for the analysis of single cell data with specific relevance to the identification of the cell (sub)types, cell phenotypes, and cell-cell signaling present in a tissue. Improved support for these single cell analysis tasks will have a meaningful impact on the basic research supporting precision medicine applications, especially in the immunology domain.

## Acknowledgments

This work was funded by National Institutes of Health grants R35GM146586, R21CA253408, P20GM130454 and P30CA023108. We would like to acknowledge the supportive environment at the Geisel School of Medicine at Dartmouth where this research was performed.

